# Genome sequencing of the perciform fish *Larimichthys crocea* provides insights into stress adaptation

**DOI:** 10.1101/008136

**Authors:** Jingqun Ao, Yinnan Mu, Li-Xin Xiang, DingDing Fan, MingJi Feng, Shicui Zhang, Qiong Shi, Lv-Yun Zhu, Ting Li, Yang Ding, Li Nie, Qiuhua Li, Wei-ren Dong, Liang Jiang, Bing Sun, XinHui Zhang, Mingyu Li, Hai-Qi Zhang, ShangBo Xie, YaBing Zhu, XuanTing Jiang, Xianhui Wang, Pengfei Mu, Wei Chen, Zhen Yue, Zhuo Wang, Jun Wang, Jian-Zhong Shao, Xinhua Chen

## Abstract

The large yellow croaker *Larimichthys crocea* (*L. crocea*) is one of the most economically important marine fish in China and East Asian countries. It also exhibits peculiar behavioral and physiological characteristics, especially sensitive to various environmental stresses, such as hypoxia and air exposure. These traits may render *L. crocea* a good model for investigating the response mechanisms to environmental stress. To understand the molecular and genetic mechanisms underlying the adaptation and response of *L. crocea* to environmental stress, we sequenced and assembled the genome of *L. crocea* using a bacterial artificial chromosome and whole-genome shotgun hierarchical strategy. The final genome assembly was 679 Mb, with a contig N50 of 63.11 kb and a scaffold N50 of 1.03 Mb, containing 25,401 protein-coding genes. Gene families underlying adaptive behaviours, such as vision-related crystallins, olfactory receptors, and auditory sense-related genes, were significantly expanded in the genome of *L. crocea* relative to those of other vertebrates. Transcriptome analyses of the hypoxia-exposed *L. crocea* brain revealed new aspects of neuro-endocrine-immune/metabolism regulatory networks that may help the fish to avoid cerebral inflammatory injury and maintain energy balance under hypoxia. Proteomics data demonstrate that skin mucus of the air-exposed *L. crocea* had a complex composition, with an unexpectedly high number of proteins (3,209), suggesting its multiple protective mechanisms involved in antioxidant functions, oxygen transport, immune defence, and osmotic and ionic regulation. Our results provide novel insights into the mechanisms of fish adaptation and response to hypoxia and air exposure.

## Introduction

Teleost fish, nearly half of all living vertebrates, display an amazing level of diversity in body forms, behaviors, physiologies, and environments that they occupy. Strategies for coping with diverse environmental stresses have evolved in different teleost species. Therefore, teleost fish are considered to be good models for investigating the adaptation and response to many natural and anthropogenic environmental stressors (Gracey et al. 2001; Cossins and Crawford 2005; van der Meer et al. 2005). Recent genome-sequencing projects in several fish have provided insights into the molecular and genetic mechanisms underlying their responses to some environmental stressors (Star et al. 2011; Schartl et al. 2013; Chen et al. 2014). However, to better clarify the conserved and differentiated features of the adaptive response to specific stresses and to trace the evolutionary process of environmental adaptation and response in teleost fish, insight from more teleost species with different evolutionary positions, such as Perciformes, is required. Perciformes are by far the largest and most diverse order of vertebrates, and thus offer a large number of models of adaptation and response to various environmental stresses.

The large yellow croaker, *Larimichthys crocea* (*L. crocea*), is a temperate-water migratory fish that belongs to the order Perciformes and the family Sciaenidae. It is mainly distributed in the southern Yellow Sea, the East China Sea, and the northern South China Sea. *L. crocea* is one of the most economically important marine fish in China and East Asian countries due to its rich nutrients and trace elements, especially selenium. In China, the annual yield from *L. crocea* aquaculture exceeds that of any other net-cage-farmed marine fish species (Liu et al. 2013; Liu et al. 2014). Recently, the basic studies on genetic improvement for growth and disease resistance traits of *L. crocea* are increasingly performed for farming purpose (Ning et al. 2007; Mu et al. 2010; Liu et al. 2013; Ye et al. 2014). *L. crocea* also exhibits peculiar behavioral and physiological characteristics, such as loud sound production, high sensitivity to sound, and well-developed photosensitive and olfactory systems (Su 2004; Zhou et al. 2011). Most importantly, *L. crocea* is especially sensitive to various environmental stresses, such as hypoxia and air exposure. For example, the response of its brain to hypoxia is quick and robust, and a large amount of mucus is secreted from its skin when it is exposed to air (Su 2004; Gu and Xu 2011). These traits may render *L. crocea* a good model for investigating the response mechanisms to environmental stress. Several studies have reported transcriptomic and proteomic responses of *L. crocea* to pathogenic infections or immune stimuli (Mu et al. 2010; Yu et al. 2010; Mu et al. 2014). The effect of hypoxia on the blood physiology of *L. crocea* has been evaluated (Gu and Xu 2011). However, little is known about the molecular response mechanisms of *L. crocea* against environmental stress.

To understand the molecular and genetic mechanisms underlying the responses of *L. crocea* to environmental stress, we sequenced its whole genome. Furthermore, we sequenced the transcriptome of the hypoxia-exposed *L. crocea* brain and profiled the proteome of its skin mucus under exposure to air. Our results revealed the molecular and genetic basis of fish adaptation and response to hypoxia and air exposure.

### Results

#### Genome features

We applied a bacterial artificial chromosome (BAC) and whole-genome shotgun (WGS) hierarchical assembly strategy for the *L. crocea* genome to overcome the high levels of genome heterozygosity (Table 1; **Supplemental Fig. S1-S2**). The 42,528 BACs were sequenced by the HiSeq 2000 platform and each BAC was assembled by SOAPdenovo (Luo et al. 2012) (**Supplemental Table S1**). The total length of all combined BACs was 3,006 megabases (Mb), which corresponded to approximately 4.3-fold genome coverage (**Supplemental Tables S2-S3**). All BAC assemblies were then merged into super-contigs and oriented to super-scaffolds with large mate-paired libraries (2-40 kb). Gap filling was made with reads from short insert-sized libraries (170-500 bp) (**Supplemental Tables S3-S4**). In total, we sequenced 563-fold coverage bases of the estimated 691 Mb genome size. The final assembly was 679 Mb, with a contig N50 of 63.11 kb and a scaffold N50 of 1.03 Mb (Table 1). The 672 longest scaffolds (11.2% of all scaffolds) covered more than 90% of the assembly (**Supplemental Table S5**). To assess the completeness of the *L. crocea* assembly, 52-fold coverage paired-end high-quality reads were aligned against the assembly (**Supplemental Fig. S3**). More than 95.63% of the generated reads could be mapped to the assembly. Furthermore, the integrity of the assembly was validated by the successful mapping of 98.80% of the transcripts from the mixed-tissue transcriptomes (**Supplemental Table S6**). These results indicate that the genome assembly of *L. crocea* has high coverage and is of high quality (**Supplemental Table S7**).

**Table 1.** Summary of the *Larimichthys crocea* genome

| Sequencing | Insert Size (bp) | Total Data (Gbp) | Coverage (×) |
| --- | --- | --- | --- |
| BAC | 120 k | 324.73 | 469.94 |
| WGS | 170–500 | 36.22 | 52.42 |
|  | 2 k–40 k | 34.26 | 49.58 |
| Assembly | N50 (bp) | Longest (kbp) | Size (Mbp) |
| Contig | 63.11 k | 716.89 | 661.33 |
| Scaffold | 1.03 M | 4,914.79 | 678.96 |
| Annotation | Number | Total Length (Mbp) | Percentage Of Genome (%) |
| Repeat | 1,454,906 | 122.92 | 18.10 |
| Gene | 25,401 | 350.95 | 51.69 |
| Exon | 251,617 | 44.86 | 6.61 |

BAC = bacterial artificial chromosome; WGS = whole genome shotgun.

The repetitive elements comprise 18.1% of the *L.crocea* genome (**Supplemental Table S8**), which is a relatively low percentage when compared with other fish species, such as *Danio rerio* (52.2%), *Gadus morhua* (25.4%), and *Gasterosteus aculeatus* (25.2%). This suggests that *L. crocea* may have a more compact genome (**Supplemental Tables S9-S10**). We identified 25,401 protein-coding genes based on *ab initio* gene prediction and evidence-based searches from the reference proteomes of six other teleost fish and humans **(Supplemental Fig. S4; Table S11)**, in which 24,941 genes (98.20% of the whole gene set) were supported by homology or RNAseq evidence (**Supplemental Fig. S5**). Over 97.35% of the inferred proteins matched entries in the InterPro, SWISS-PROT, KEGG or TrEMBL database (**Supplemental Table S12**).

#### Phylogenetic relationships and genomic comparison

*L. crocea* is the first species of *Sciaenidae* of the order *Perciformes* with a complete genome available, therefore we estimated its phylogenetic relationships to seven other sequenced teleost species based on 2,257 one-to-one high-quality orthologues, using the maximum likelihood method. According to the phylogeny and the fossil record of teleosts, we dated the divergence of *L. crocea* from the other teleost species to approximately 64.7 million years ago (Fig. 1A). We also detected 19,283 orthologous gene families (**Supplemental Table S3**), of which 14,698 families were found in *L. crocea*. The gene components of *L. crocea* were similar to those of *D. rerio* (Fig. 1B). The gene contents in four representative teleost species and *L. crocea* genomes were also analysed, and 11,205 (76.23%) gene families were found to be shared by five teleosts (Fig. 1C). We confirmed that the one-to-one orthologous genes of *G. aculeatus* and *L. crocea* have higher sequence identities from the distribution of the percent identity of proteins (Fig. 1D), which indicates that Sciaenidae has a closer affinity to Gasterosteiformes and coincides with our genome-level phylogeny position.

**Figure 1.**
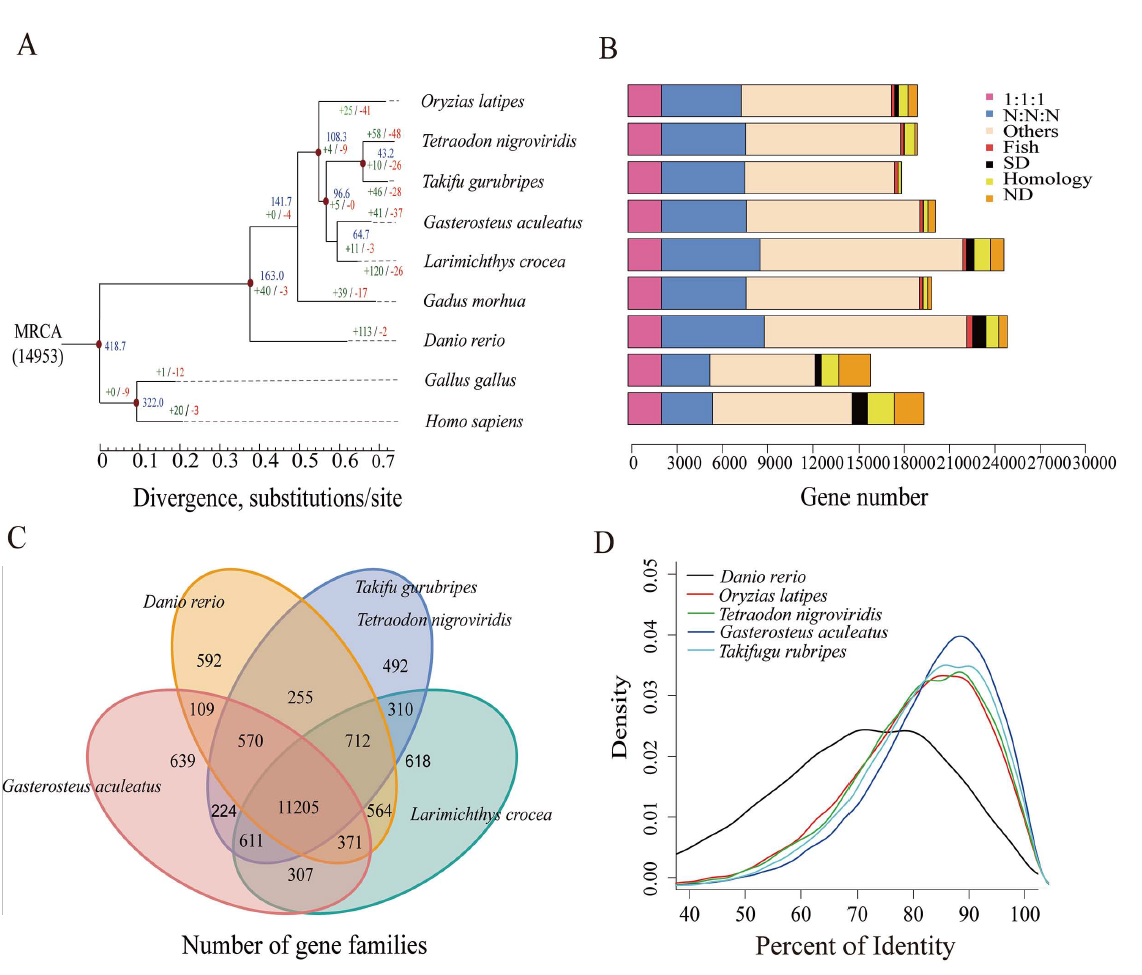
Phylogenetic tree of and orthologous genes in *L. crocea* and other vertebrates. (**A**) The phylogenetic tree was constructed from 2,257 single-copy genes with 3.18 M reliable sites by maximum likelihood methods. The red points on six of the internal nodes indicate fossil calibration times in the analysis. Blue numbers indicate the divergence time (Myr, million years ago), and the green and red numbers represent the expanded and extracted gene families, respectively, in *L. crocea*. (**B**) The different types of orthologous relationships are shown. “1:1:1” = universal single-copy genes; “N:N:N” = orthologues exist in all genomes; “Fish” = fish-specific genes; “SD” = genes that have undergone species-specific duplication; “Homology” = genes with an e-value less than 1e-5 by BLAST but do not cluster to a gene family; “ND” = species-specific genes; and “Others” = orthologues that do not fit into the other categories. (**C**) The shared and unique gene families in five teleost fish are shown in the Venn diagram. (**D**) Distribution of the identity values of orthologous genes is compared among *L. crocea* and other teleosts.

Furthermore, 121 significantly expanded and 27 contracted gene families (*P* < 0.01) were identified by comparing the family size of *L. crocea* with that of the other vertebrates used in the phylogenetic analysis (**Supplemental Tables S14-S15**). Based on the ratios of the number of nonsynonymous substitutions per nonsynonymous site (K_a_) to the number of synonymous substitutions per synonymous site (K_s_; K_a_/K_s_ ratios) in a branch-site model of PAML (Yang 1997), 92 genes were found to be positively selected in *L. crocea* compared with their orthologues in the other six teleost species (*P* < 0.001, **Supplemental Table S16**).

#### Unique genetic features of the *L. crocea*

*L. crocea* is a migratory fish with good photosensitivity, olfactory detection, and sound perception, and it contains high levels of selenium (Su 2004). Our genomic analyses provide genetic basis for these behavioral and physiological characteristics. Several crystallin genes (*crygm2b*, *cryba1*, and *crybb3*), which encode proteins that maintain the transparency and refractive index of the lens (Chen et al. 2014), were significantly expanded in the genome of *L. crocea* relative to those of other sequenced teleosts (**Supplemental Table S17**). Phylogenetic analysis showed that the crystallin genes from *L. crocea* cluster together, indicating that these genes were specifically duplicated in *L. crocea* lineage (**Supplemental Fig. S6**). The specific expansion of these crystallin genes may be helpful for improving photosensitivity by increasing lens transparency, thereby enabling the fish to easily find food and avoid predation underwater.

We also identified 112 olfactory receptor (OR)-like genes from the *L. crocea* genome (**Supplemental Table S18**; **Fig. S7**), and almost all of them (111) have been reported to be expressed in the olfactory epithelial tissues of *L. crocea* (Zhou et al. 2011). The majority of these genes (66) were classified into the “delta” group, which is important for the perception of water-borne odorants (Niimura 2009). *L. crocea* also possessed the highest number of genes that were classified into the “eta” group (30, *P* < 0.001), and these genes may contribute to the olfactory detection abilities, which could be useful for feeding and migration (Li et al. 1995).

*L. crocea* is named for its ability to generate strong repetitive drumming sounds, especially during reproduction (Su 2004). For good communication, fish have developed high sensitivities to environmental sound. Three important auditory genes, otoferlin (*OTOF*), *claudinj*, and otolin 1 (*OTOL1*), were significantly expanded in the *L. crocea* genome (*P* < 0.01, **Supplemental Table S19**). These expansions may contribute to the detection of sound signaling during communication, and thus to reproduction and survival (Eisen and Ryugo 2007).

Selenium is highly enriched in *L. crocea* (Su 2004), and it is mainly present as selenoproteins. We used the SelGenAmic-based selenoprotein prediction method (Jiang et al. 2010) to analyse the *L. crocea* genome and identified 40 selenoprotein genes, which is the highest number among all sequenced vertebrates (**Supplemental Table S20**). Interestingly, five copies of *MsrB1*, which encodes methionine sulfoxide reductase, were found in *L. crocea* (*MsrB1a, MsrB1b, MsrB1c*, *MsrB1d*, and *MsrB1e*), whereas only two copies (*MsrB1a* and *MsrB1b*) were found in other fish, thus suggesting its broader specificity to reduce all possible substrates (Vandermarliere et al. 2014).

#### Characterization of the *L. crocea* immune system

Approximately 2,524 immune-relevant genes were annotated in the *L. crocea* genome, including 819 innate immune-relevant genes and 1,705 adaptive immune-relevant genes (**Supplemental Table S21**). *L. crocea* has a relatively complete innate immune system, whereas its adaptive immune system may possess unique characteristics. The CD8^+^ T and CD4^+^ T-helper type 1 (Th1)-type immune systems are well conserved in *L. crocea*, and almost all CD8^+^ T and CD4^+^ Th1 cell-related genes were found (Fig. 2A). Moreover, the genes related to Th17 cell-and γδ-T cell-mediated mucosal immune responses were conserved in *L. crocea*. These observations suggest that *L. crocea* may exhibit powerful cellular and mucosal immunity. However, the CD4^+^ Th2-type immunity seemed to be weak in *L. crocea*, as suggested by the absence of many CD4^+^ Th2-related genes and humoral immune effectors (Fig. 2A). We detected gene expansions in several of these immune-relevant genes, including those encoding lectin receptors (*CLEC17A*), a classical complement component (*C1q*), apoptosis regulator (*BAX*), and immunoglobulins (*IgHV*) (*P* < 0.01, **Supplemental Table S22**). Expansions were also observed in the genes encoding four key proteins for mammalian antiviral immunity: tripartite motif containing 25 (TRIM25), cyclic GMP-AMP synthase (cGAS), DDX41, and NOD-like receptor family CARD domain containing 3 (NLRC3) (Fig. 2B). However, retinoic acid-inducible gene-1 (*RIG-I*), which initiates antiviral signaling pathway in mammals, was not found in the *L. crocea* genome and transcriptome (Mu et al. 2010; Mu et al. 2014). The teleost *RIG-I* has been identified only in limited fish species, such as cyprinids and salmonids, and its absence suggests that it may have been lost from particular fish genomes (Hansen et al. 2011). Furthermore, laboratory of genetics and physiology 2 (LGP2) can serve as a suppressor to block RIG-I-and melanoma differentiation-associated protein 5 (MDA5)-elicited signaling in mammals, but LGP2 in fish is able to bind to poly(I:C) to trigger interferon production (Chang et al. 2011), thereby acting as a substitute for RIG-I (Fig. 2B). The expanded *TRIM25* (54 copies, **Supplemental Fig. S8**) may trigger the ubiquitination of interferon-β promoter stimulator-1 (IPS-1), thus allowing for interferon regulatory factor 3 (IRF3) phosphorylation and antiviral signaling initiation (Castanier et al. 2012). *DDX41* and *cGAS* encode intracellular DNA sensors, which can activate stimulator of interferon genes (STING) and TANK-binding kinase 1 (TBK1) to induce type I interferons (Zhang et al. 2011; Gao et al. 2013). *L. crocea* contained 76 copies of *NLRC3* (**Supplemental Fig. S9**), which encodes regulators that prevent type I interferon overproduction (Zhang et al. 2014). The expansions of these virus-response genes suggest their enhanced roles in innate antiviral immunity, which may explain why *L. crocea* is less susceptible to viral infection.

**Figure 2.**
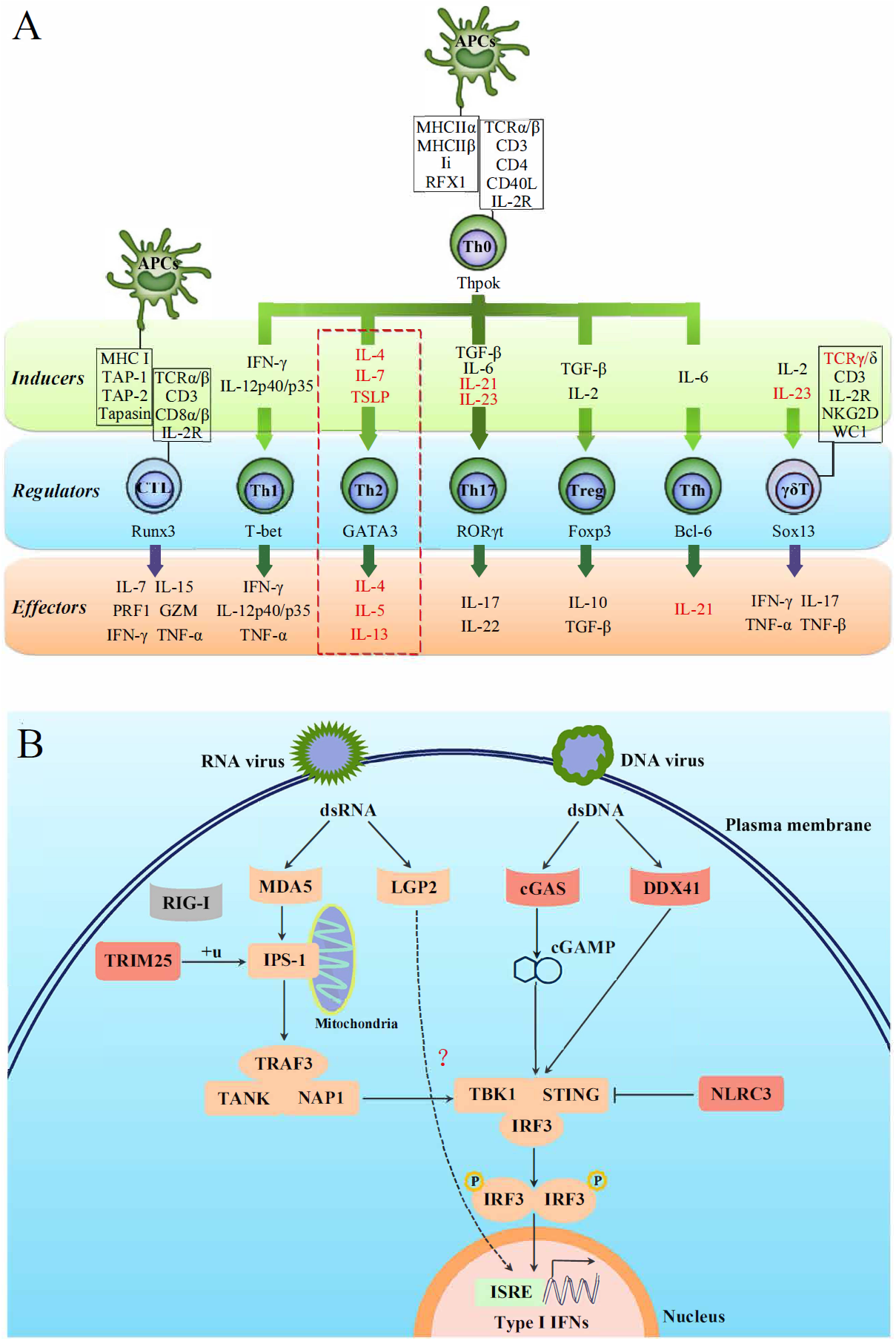
Characterisation of the T-cell lineages in *L. crocea* adaptive immunity and the expanded genes in antiviral immunity. (**A**) A schematic diagram summarising genes related to different T-cell lineages in *L. crocea* is shown. The inducible factors, the main regulatory transcriptional factors, and the immune effectors of T cells are present in green, blue, and orange backgrounds, respectively. The genes that have been annotated by genome survey are shown in black, and the unannotated genes are shown in red. The dashed square outlines the possible incomplete Th2 cell-mediated humoral immunity of *L. crocea*. (**B**) Several key genes are expanded in the antiviral immunity pathways in *L. crocea*. The genes that have been identified in the *L. crocea* genome are shown in orange boxes, and the lost gene (*RIG-I*) is shown in the grey box. LGP2 is able to bind to double-stranded RNA (dsRNA) to trigger interferon production, but the adaptor molecule of LGP2 is still unknown in fish. The red boxes indicate gene families (*TRIM25, cGAS, DDX41*, and *NLRC3*) that are expanded in *L. crocea.* The arrow represents induction, and the interrupted line represents inhibition.

#### Stress response under hypoxia

The brain allows rapid and coordinated responses to the environmental stress by driving the secretion of hormones. Therefore, we studied the response of the *L. crocea* brain to hypoxia. We sequenced seven transcriptomes of the brains at different times of hypoxia exposure and found that 8,402 genes were differentially expressed at one or more time points (false discovery rate [FDR] ≤ 0.001, fold change > 2; **Supplemental Fig. S10**). Hypoxia stress induced a response with the largest number of genes (4,535 genes) at 6 h (**Supplemental Fig. S11**), indicating that genes with regulated expression at 6 h may be critical for the response. Hypoxia stress can induce the response of the central neuroimmune system, in which brain neuropeptides, endocrine hormones, and inflammatory cytokines closely participate (Herman and Cullinan 1997; Yang et al. 2012; Lemos Vde et al. 2013). However, the precise regulatory networks among these factors have not yet been fully delineated. Our transcriptome analysis of *L. crocea* brains under hypoxia stress may outline a novel hypothalamic-pituitary-adrenal (HPA) axis-endothelin-1 (ET-1)/adrenomedullin (ADM)-interleukin (IL)-6/tumor necrosis factor (TNF)-α feedback regulatory loop that is involved in the neuro-endocrine-immune network during hypoxia responses (Fig. 3; **Supplemental Table S23; Fig. S12**).

**Figure 3.**
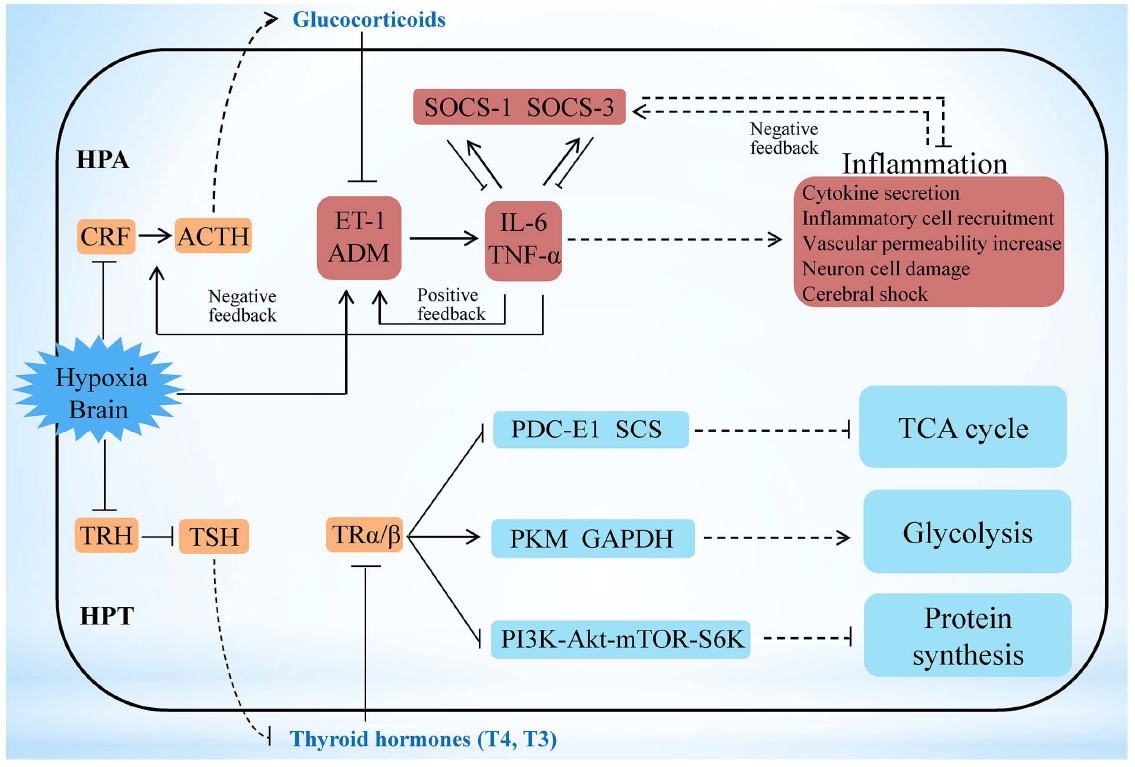
Hypoxia stress exerts responses involving the HPA and HPT axes. Under hypoxia, a potential neuro-endocrine-immune/metabolism network contributes to the regulation of moderate inflammation and the maintenance of energy balance. Hypoxia can initially promote *ET-1* and *ADM* expression, after which it increases pivotal inflammatory cytokines, such as IL-6 and TNF-α in the brain, to induce cerebral inflammation. ET-1/ADM and IL-6/TNF-α form a positive feedback loop to amplify cerebral inflammation. Afterwards, the hypothalamic-pituitary-adrenal (HPA) axis-glucocorticoids pathway and SOCS family members (SOCS-1 and SOCS-3) can inhibit *IL-6/TNF-α* expression, which constitutes the negative feedback loops with IL-6/TNF-α to modify cerebral inflammation. The HPT axis was inhibited in *L. crocea* brains during the early period of hypoxia, thus leading to a decrease in thyroid hormone production. Thyroid hormones subsequently inhibited ribosomal biogenesis and protein translation by the PI3K-Akt-mTOR-S6 K signaling pathway. Down-regulation of HPT axis-thyroid hormones also repressed the tricarboxylic acid (TCA) cycle and accelerated the anaerobic glycolytic pathway in the brain, along with increases in the exposure to hypoxia. Genes related to the neuro-endocrine system (orange), immunity (red), and metabolic system and protein synthesis (blue) are indicated. The outer border indicates the brain of *L. crocea*. The arrow represents promotion, and the interrupted line represents inhibition. Solid lines indicate direct relationships between genes. Dashed lines indicate that more than one step is involved in the process.

Results from transcriptome analyses show that the key HPA axis-relevant genes (corticotropin-releasing factor [*CRF*], CRF receptor 1 [*CRFR1*], pro-opiomelanocortin [*POMC*], and CRF-binding protein [*CRFBP*]) in the *L. crocea* brain displayed a down-up-down-up (W-type) dynamic expression pattern under hypoxia stress (**Supplemental Fig. S12)**. The HPA axis can strictly control the production of glucocorticoids (Nadeau and Rivest 2003; Sorrells and Sapolsky 2007), and glucocorticoids are suppressors of ET-1 and ADM, which are both involved in cerebral inflammation in mammals (Takahashi et al. 2003; Hayashi et al. 2004). Meanwhile, the dynamic expression levels of *ET-1* and *ADM* clearly showed a typical M-type pattern (up-down-up-down), and the time of inflexion point corresponded with that of *CRF*, *CRFR1*, *POMC*, and *CRFBP*. These observations suggest the existence of a feedback regulatory pathway between the HPA axis and ET-1/ADM under hypoxia stimulation. Notably, the expression of *IL-6/TNF-α* showed the M-type pattern and was consistent with that of *ET-1/ADM* (**Supplemental Fig. S12)**. These coordinated and fluctuating expression patterns indicate that hypoxia may induce the expression of *ET-1/ADM* and *IL-6/TNF-α* and trigger a positive feedback loop between them (Fig. 3). Furthermore, ET-1/ADM-IL-6/TNF-α may activate the HPA axis, and the latter subsequently induces glucocorticoids and generates a negative feedback to inhibit *ET-1/ADM* and *IL-6/TNF-α* expression to reduce inflammatory response in brain. This suggestion could be supported by previous reports in mammals (Mastorakos et al. 1993; Kitamuro et al. 2000; Earley et al. 2002; Takahashi et al. 2003).

*L. crocea* also exhibits other protective mechanisms, such as the suppressors of cytokine signaling (SOCS)-dependant regulatory mechanism, to avoid inflammation-induced cerebral injury. Both *SOCS-1* and *SOCS-3* in the *L. crocea* brain display opposite expression patterns against *IL-6* and *TNF-α* (**Supplemental Fig. S12)**. Thus, SOCS-1 and SOCS-3 may have complementary roles in down-regulating *IL-6* and *TNF-α*, and both IL-6 and TNF-α have reciprocal functions to induce *SOCS-1* and *SOCS-3* expression (Fig. 3). These results suggest that a SOCS-1/3-dependent feedback regulation may exist in the process against hypoxia-induced cerebral inflammation in *L. crocea*.

Hypoxia can influence the hypothalamic-pituitary-thyroid (HPT) axis (Hou and Du 2005). HPT axis was found to regulate protein synthesis and glucose metabolism by production of thyroid hormones (Yen 2001). Here, the major HPT axis-related genes (thyrotropin-releasing hormone [*TRH*], TRH receptor [*TRHR*], thyroid-stimulating hormone [*TSH*], and TSH receptor [*TSHR*]) were significantly down-regulated in the *L. crocea* brain at 1 h to 6 h under hypoxia (**Supplemental Table S24**), thus indicating that the HPT axis may be inhibited during the early period of hypoxia. Inhibition of the HPT axis leads to a decrease in the production of thyroid hormones. Furthermore, thyroid hormones can regulate ribosomal biogenesis and protein translation by the PI3K-Akt-mTOR-S6 K signaling pathway (Kenessey and Ojamaa 2006). In this study, the mRNA levels of *PI3K*, *S6K*, and most of the components of the protein translation machinery, including the ribosomal proteins and eukaryotic translation initiation factors (*eIF-1, −2, −3, −5* and *-6*), were all down-regulated under hypoxia (**Supplemental Table S25**). This suggests that the HPT axis may inhibit protein synthesis under hypoxia by decreasing the production of thyroid hormones (Fig. 3), which is beneficial for saving energy during hypoxia stress. Thyroid hormones can also accelerate the oxidative metabolism of glucose and inhibit the glycolytic anaerobic pathway (Sabell et al. 1985). Our transcriptome analyses show that genes involved in the tricarboxylic acid (TCA) cycle (pyruvate dehydrogenase complex [*PDC-E1*], succinyl-CoA synthetase [*SCS*], and fumarate hydratase [*FH*]) were down-regulated 12 h later under hypoxia, whereas glycolysis-related genes, such as pyruvate kinase (*PKM*), glyceraldehyde 3-phosphate dehydrogenase (*GAPDH*), *GPI*, and aldolase A (*ALDOA*), were greatly increased at 1 h (280-, 130-, 73- and 12-fold, respectively) (**Supplemental Table S24**). The down-regulation of HPT axis-thyroid hormones may inhibit the TCA cycle and accelerate the anaerobic glycolytic pathway in the brain during hypoxia exposure (Fig. 3). The repression of the TCA cycle and the strong induction of the anaerobic glycolytic pathway resulted in a physiological shift from aerobic to anaerobic metabolism, where fish utilise O_2_-independent mechanisms to produce adenosine triphosphate (ATP). However, the mRNA levels of hypoxia-inducible factor (HIF)-1α, which are significantly up-regulated under hypoxia in mammals (Dayan et al. 2006; Benita et al. 2009), were not significantly changed in the *L. crocea* brain (**Supplemental Table S24**). It is possible that the HIF-1α-mediated mechanism may not be essential for the hypoxia response in the *L. crocea* brain during the early period of hypoxia. These results suggest that the HPT axis-mediated effects may play major roles in response to hypoxia by reorganizing energy consumption and energy generation.

#### Mucus components and function

The skin mucus is considered as the first defensive barrier between fish and its aquatic environment, and it plays a role in a number of functions, including locomotion, antioxidant responses, respiration, disease resistance, communication, ionic and osmotic regulation (Shephard 1994). However, the exact mechanisms underlying these functions remain unknown. Mucus is composed mainly of the gel-forming macromolecule mucin and water (Subramanian et al. 2008). We identified 159 genes that are implicated in mucin biosynthesis and mucus production in the *L. crocea* genome (**Supplemental Table S26**), based on previous studies in mammals (Pluta et al. 2012). This indicates that the mucin synthetic pathway is conserved between fish and mammals. Among these gene families, GALNT, which encodes N-acetylgalactosaminyl transferases (Guzman-Aranguez et al. 2009), was significantly expanded in *L. crocea* (27 copies versus 15–20 copies in other fish) (**Supplemental Fig. S13**). Syntaxin-11 was also expanded. Additionally, genes encoding syntaxin-binding protein 1 and syntaxin-binding protein 5, which are related to mucus secretion, were positively selected in the *L. crocea* genome (**Supplemental Table S16**). The expansion and positive selection of these genes may explain why the *L. crocea* secretes more mucus than other fish under stress.

We identified 22,054 peptides belonging to 3,209 genes in the *L. crocea* skin mucus proteome, and this accounted for more than 12% of the protein-coding genes in the genome (**Supplemental Table S27**). The complexity of the *L. crocea* mucus presumably relates to the multitude of its biological functions that allow the fish to survive and adapt to environmental changes. The over-represented functional categories were oxidoreductase activity (GO:0016491, *P* = 1.58×10^−35^, 223 proteins), peroxidase activity (GO:0004601, *P* = 0.0075, nine proteins), oxygen binding (GO:0019825, *P* = 0.0011, eight proteins), and ion binding (GO:0043167, *P* = 2.21×10^−6^, 347 proteins) (Fig. 4A; **Supplemental Fig. S14**). Two hundred and thirty-two antioxidant proteins that were related to oxidoreductase activity and peroxidase activity were highly enriched in the *L. crocea* mucus, and they included peroxiredoxins, glutathione peroxidase, and thioredoxin (**Supplemental Table S28**). These proteins intercept and degrade environmental peroxyl and hydroxyl radicals from aqueous environments (Cross et al. 1984). Therefore, the presence of high-abundance antioxidant proteins in the skin mucus may have the potential to protect fish against air exposure-induced oxidative damage (Fig. 4B). Eight proteins related to oxygen transport, including hemoglobin subunits α1, αΑ, αD, β, and β1, and cytoglobin-1, were identified in the *L. crocea* skin mucus (**Supplemental Table S29**). The abundant expression of hemoglobin may contribute to the binding and holding of oxygen for respiration. Various immune molecules that provide immediate protection to fish from potential pathogens, such as lectins, lysozymes, C-reactive proteins, complement components, immunoglobulins, and chemokines, were also found in the *L. crocea* skin mucus (**Supplemental Table S30**). To date, the mechanisms of osmotic and ionic regulation of the skin mucus have not been confirmed (Shephard 1994). In this study, a large number of ion-binding proteins were identified in the *L. crocea* mucus (**Supplemental Table S31**). These proteins and the layer of mucus may have a role in limiting the diffusion of ions on the surface of the fish (Fig. 4B). However, a substantial proportion of the proteins, which are highly present in the skin mucus of fish under air exposure, play an unknown role in the mucus response.

**Figure 4.**
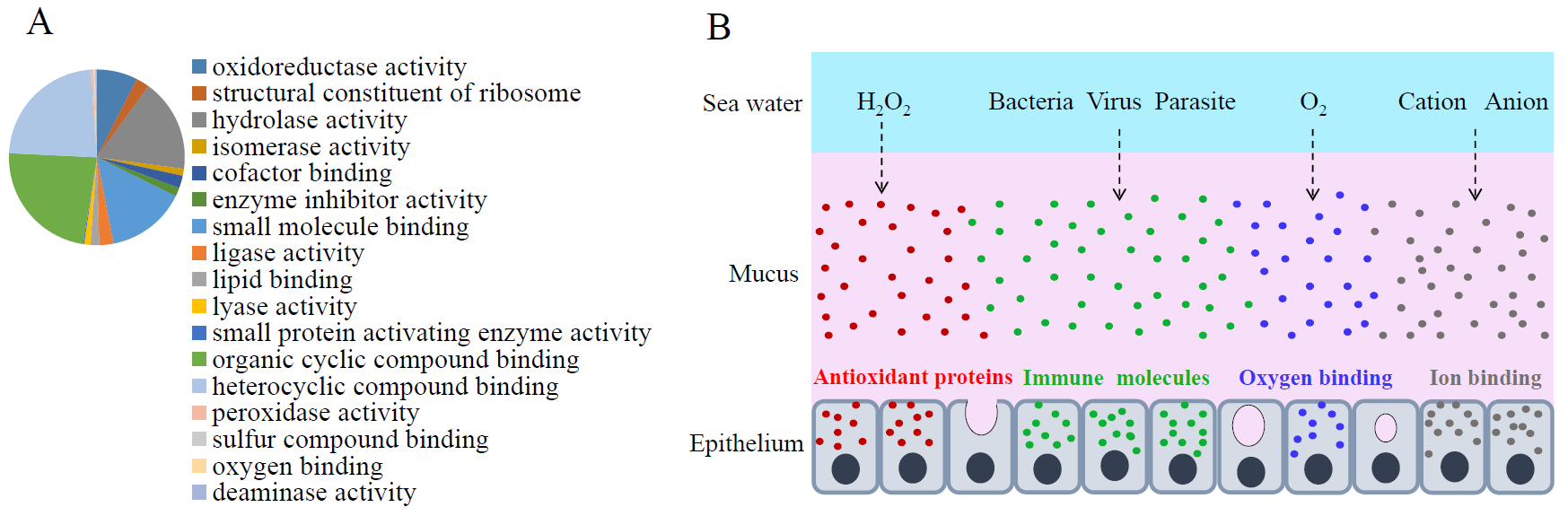
Skin mucus proteins are overexpressed in air-exposed *L. crocea*. (**A**) The distribution of mucus proteins in the molecular function class of Gene Ontology is shown. The over-represented functional categories are indicated in the pie chart. (**B**) A representation of the functional mechanisms of the mucus barrier is shown. The continuously replenished thick mucus layer can retain a large number of antioxidant, immune, oxygen-binding, and ion-binding molecules, which are involved in antioxidant functions, immune defence, oxygen transport, and osmotic and ionic regulation, respectively.

### Discussion

We sequenced and assembled the genome of the large yellow croakerr (*L. crocea*) using BACs and the WGS hierarchical assembly strategy. This methodology is effective for high-polymorphism genomes and produces a high quality genome assembly, with the 63.11 kb contig N50 and 1.03 Mb scaffold N50 (Table 1). Support from the 563-fold coverage of genome yields high single-base resolution and 98.80% completeness of the coding region (**Supplemental Table S6**). Further genomic analyses showed the significant expansion of several gene families, such as vision-related crystallins, olfactory receptors, and auditory sense-related genes, and provided a genetic basis for the peculiar behavioral and physiological characteristics of *L. crocea*.

During the early stages of hypoxia stress, the induction of ET-1/ADM and IL-6/TNF-α generates the primary protective effect to increase blood pressure, enhance vascular permeability and trigger inflammatory response (Bona et al. 1999; Taylor et al. 2005). These mechanisms maintain the brain oxygen supply and resist pathogen infection when the blood brain barrier is disrupted by hypoxia (Kaur and Ling 2008). As the stress response progresses, several natural brakes, including HPA axis-Glucocorticoids and SOCS family members, exhibit secondary protection effects to avoid excessive inflammatory responses in the brain. Our transcriptome results show that a novel HPA axis-ET-1/ADM-IL-6/TNF-α feedback regulatory loop in neuro-endocrine-immune networks contributed to the protective effect and regulated moderate inflammation under hypoxia stress (Fig. 3). On the other hand, the hypoxia-induced down-regulation of the HPT axis may lead to the inhibition of protein synthesis and the activation of anaerobic metabolism (Fig. 3; **Supplemental Tables S24-S25**). Inhibition of protein synthesis principally contributes to the reduction in cellular energy consumption during hypoxia (Gracey et al. 2001; Richards 2011). Activation of anaerobic metabolism facilitates O_2_-independent ATP production under hypoxia, albeit with low ATP yield (Richards 2011). Therefore the reduction in ATP consumption through the HPT axis-mediated inhibition of protein synthesis matched the lower ATP yield by the HPT axis-activated anaerobic metabolism, which may aid to maintain cellular energy balance under hypoxia, thus extending fish survival. Hence, our results reveal new aspects of neuro-endocrine-immune/metabolism regulatory networks that may help the fish to avoid cerebral inflammatory injury and maintain energy balance under hypoxia stress. These discoveries will help to improve current understanding of neuro-endocrine-immune/metabolism regulatory networks and protective mechanisms against hypoxia-induced cerebral injury in vertebrates, providing clues for research on the pathogenesis and treatment of hypoxia-induced cerebral diseases.

Amazingly, 3,209 different proteins were identified in the *L. crocea* skin mucus under air exposure. Of these, oxidoreductase activity-, oxygen binding-, immunity-, and ion binding-related proteins were enriched (Fig. 4A; **Supplemental Fig. S14**). The increase in secretion of the skin mucus of *L. crocea* under air exposure may reflect a physiological adjustment of the fish to cope with environmental changes, and the complex components suggest that the skin mucus exerts multiple protective mechanisms, which are involved in antioxidant functions, oxygen transport, immune defence, and osmotic and ionic regulation (Fig. 4B). These results expand our knowledge of skin mucus secretion and function in fish, highlighting its importance in response to stress. In addition, the mucus proteome shares many proteins with the mucus from humans and other animals (Lee et al. 2011; Rodriguez-Pineiro et al. 2013). These characteristics thus make *L. crocea* a pertinent model for studying mucus biology.

In summary, our sequencing of the genome of the large yellow croaker provided the genetic basis for its peculiar behavioral and physiological characteristics. Results from transcriptome analyses revealed new aspects of neuro-endocrine-immune/metabolism regulatory networks that may help the fish to avoid cerebral inflammatory injury and maintain energy balance under hypoxia stress. Proteomic profiling suggested that the skin mucus of the fish exhibits multiple protective mechanisms in response to air-exposure stress. Overall, our results revealed the molecular and genetic basis of fish adaptation and response to hypoxia and air exposure. In addition, the data generated by this study will facilitate the genetic dissection of aquaculture traits in this species and provide valuable resources for the genetic improvement of the meat quality and production of *L. crocea*.

### Materials and Methods

#### Genome assembly annotation

The wild *L. crocea* individuals were collected from the Sanduao sea area in Ningde, Fujian, China. Genomic DNA was isolated from the blood of a female fish by using standard molecular biology techniques for BAC library construction and sequencing by the HiSeq 2000 Sequencing System in BGI (Beijing Genomics Institute, Shenzhen, China). Subsequently, low-quality and duplicated reads were filtered out, and sequencing errors were removed. The BACs of *L. crocea* were assembled by using SOAPdenovo2 (Li et al. 2009) (http://soap.genomics.org.cn) with k-mers that ranged from 25 to 63 in size. Then, we selected the assembly with the longest scaffold N50 for gap filling. The BACs were merged together based on the overlap found by BLAT, using the custom script: Rabbit (ftp://ftp.genomics.org.cn/pub/Plutellaxylostella/Rabbit_linux-2.6.18-194.blc.tar.gz). The redundant sequences that were produced by high polymorphisms were removed by sequence depth and shared k-mer percentage. Assembly was performed by scaffolding with mate-paired libraries (2–40 kb) using SSPACE v2 (Boetzer et al. 2011), and gap filling was made by Gapcloser (http://sourceforge.net/projects/soapdenovo2/files/GapCloser/) with small-insert libraries (170–500 bp).

#### Genome annotation

For the annotation of repetitive elements, we used a combination of homology-based and *ab initio* predictions. RepeatMasker (Smit 1996-2010) and Protein-based RepeatMasking (Smit 1996-2010) were used to search Repbase, which contains a vast amount of known transcriptional elements at the DNA and protein levels. During the process of *ab initio* prediction, RepeatScout (Price et al. 2005) was used to build the *ab initio* repeat library based on k-mer, using the fit-preferred alignment score on the *L. crocea* genome. Contamination and multi-copy genes in the library were filtered out before the RepeatScout library was used to find homologs in the genome and to categorise the found repeats by RepeatMasker (Smit 1996-2010).

Gene models were integrated based on *ab initio* predictions, homologue prediction, and transcription evidence.

#### Homology-based prediction

The protein sequences of seven species (*Danio rerio, Gasterosteus aculeatus, Oreochromis niloticus, Oryzias latipes, Takifugu rubripes, Tetraodon nigroviridis*, and *Homo sapiens*) were aligned to the *L. crocea* assembly using BLAST (E-value ≤ 1e-5), and the matches with length coverage > 30% of the homologous proteins were considered as gene-model candidates. The corresponding homologous genome sequences were then aligned against the matching proteins by using Genewise (Birney et al. 2004) to improve gene models.

#### *Ab initio* prediction

Augustus (Stanke and Morgenstern 2005), SNAP (Korf 2004), and GENESCAN (Burge and Karlin 1997) were used for the *ab initio* predictions of gene structures on the repeat-masked assembly.

#### Transcriptome-based prediction

RNAseq reads from the transcriptomes of the mixed tissues of a female and a male (eleven tissues each) were aligned to the genome assembly by Tophat (Trapnell et al. 2009), which can identify splice junctions between exons. Cufflinks (Mortazavi et al. 2008) was used to obtain transcript structures.

Homology-based, *ab initio* derived and transcript gene sets were integrated to form a comprehensive and non-redundant gene set. The overlap length of each gene was verified by different methods, and genes showing 50% overlap by at least one method were selected. To eliminate false positives (genes only supported by *ab initio* methods), novel genes with the reads per kb of gene model per million of reads (rpkm) ≤ 1 were removed.

#### Evolutionary and Comparative Analyses

To detect variations in the *L. crocea* genome, we chose nine species (*Larimichthys crocea, Gasterosteus aculeatus, Takifugu rubripes, Tetraodon nigroviridis, Oryzias latipes, Gadus morhua, Danio rerio, Gallus gallus*, and *Homo sapiens*). Proteins that were greater than 50 amino acids in size were aligned by BLAST (-p blastp-e 1e-7), and Treefam (Ruan et al. 2008) was used to construct gene families for comparison.

The 2,257 single-copy genes from the gene family analysis were aligned using MUSCLE (Edgar 2004), and alignments were concatenated as a single data set. To reduce the error topology of phylogeny by alignment inaccuracies, we used Gblock (Castresana 2000) (codon model) to remove unreliably aligned sites and gaps in the alignments. The phylogenetic tree and divergence time were calculated using the PAML 3.0 (Yang 1997) package.

Gene family expansion and contraction analyses were performed by cafe (De Bie et al. 2006). For optical, olfactory receptor, and auditory system-related genes, we downloaded the genes from Swissprot or Genebank and predicted their candidates using BLAST and Genewise to determinate copy numbers. Pseudogenes produced by frame shift were removed. Phylogenetic analysis of the expanded gene families was based on maximum likelihood methods by PAML 3.0 (Yang 1997), and the phylogenetic tree was represented by EvolView (Zhang et al. 2012b).

Amino acid sequences from six representative teleosts (*Larimichthys crocea, Gasterosteus aculeatus, Danio rerio, Oryzias latipes, Takifugu rubripes,* and *Tetraodon nigroviridis*) were aligned by BLAST (-p blastp -e 1e-5 -m 8), and reciprocal-best-BLAST-hit methods were used to define orthologous genes in six teleost fish. Because alignment errors are an important concern in molecular data analysis, we made alignments of codon sequences, which are nucleotide sequences that code for proteins, using the PRANK (Loytynoja and Goldman 2010) aligner. Positive selection was inferred, based on the branch-site K_a_/K_s_ test by codeml in the PAML 3.0 package (Yang 1997).

#### Transcriptome under hypoxia

*L. crocea* (90–100 g) individuals were purchased from a mariculture farm in Ningde, Fujian, China. The fish were maintained at 25 °C in aerated water tanks (dissolved oxygen [DO] concentration: 7.8 ± 0.5 mg/L) with a flow-through seawater supply. After 7 days of acclimation, hypoxia-exposure experiments were conducted at 25 °C using published methods (Gracey et al. 2001) by bubbling nitrogen gas into an aquarium. The desired concentration of DO was detected by using a DO meter (YSI, Canada). *L. crocea* cannot maintain the aerobic pathway at DO levels below 2.0 mg/L, and it resorts to anaerobic metabolism (Gu and Xu 2011). Therefore, at the onset of hypoxia, the oxygen content in the tank was lowered from 7.8 ± 0.5 mg/L to 1.6 ± 0.2 mg/L over a 10-min period. Brains were harvested from six fish at the 1-, 3-, 6-, 12-, 24-, and 48-h time points and frozen immediately in liquid nitrogen until RNA extraction and transcriptome analyses were performed.

Total RNA was extracted from the tissues of *L. crocea* using the guanidinium thiocyanate-phenol-chloroform extraction method (Trizol, Invitrogen, USA), according to the manufacturer’s protocol. The libraries were sequenced by using the Illumina HiSeq 2000 sequencing platform with the paired-end sequencing module (Zhang et al. 2012a). After removing low-quality reads, RNAseq reads were aligned to the *L. crocea* genome with SOAPaligner/SOAP2 (Li et al. 2009). The alignment was utilised to calculate the distribution of reads on reference genes and to perform coverage analysis. If an alignment result passed quality control (alignment ratio > 70%), we proceeded in gene expression calculations and differential expression comparisons.

#### LC–MS/MS analyses and mucus protein identification

Skin mucus was collected from six healthy *L. crocea* individuals under air exposure as previously described (Subramanian et al. 2008). Briefly, the fish were anesthetised with a sub-lethal dose of Tricaine-S (100 mg/L), and transferred gently to a sterile plastic bag for 3 min to slough off the mucus under air exposure. To exclude the cell contamination, mucus was diluted in fresh, cold phosphate-buffered saline and drop-splashed onto slides, which were then air-dried. After staining with 10% Giemsa dye (Sigma, St Louis, MO, USA) for 20 min, the mucus was observed under a Nikon microscope with a 20 × objective. No cell was observed.

Proteins were extracted from a pool of skin mucus of six fish by the trichloroacetic acid-acetone precipitation method and digested by the trypsin gold (Promega, USA). The peptides were then separated by the strong cation exchange chromatography using a Shimadzu LC-20AB HPLC Pump system (Kyoto, Japan). Data acquisition was performed with a Triple TOF 5600 System (AB SCIEX, Concord, ON) fitted with a Nanospray III source (AB SCIEX, Concord, ON). All spectra were mapped by MASCOT server version 2.3.02 against the database of the *L. crocea* genome with the parameters as follows: peptide mass tolerance 0.05 Da; fragment mass tolerance 0.1 Da; fixed modifications “Carbamidomethyl (C)”; and variable modifications “Gln->pyro-Glu (N-term Q), Oxidation (M), Deamidated (NQ)”. For further analyses of the function of the mucus proteome, we selected proteins with more than two unique peptides.

### Data access

The large yellow croaker whole-genome sequence has been deposited at the DNA Data Bank of Japan (DDBJ), the European Molecular Biology Laboratory (EMBL) nucleotide sequencing database and GenBank under the same accession XXX (The data have been submitted and we are waiting for return of the accession numbers). All short-read data of WGS and BAC have been deposited in the Short Read Archive (SRA) under accession SRA159210 and SRA159209 respectively. Raw sequencing data for the transcriptome have been deposited in the Gene Expression Omnibus (GEO) under accession GSE57608.

## Acknowledgements

This work was supported by the Nation ‘863’ Project (2012AA092202); National Basic Research Program of China (2012CB114402); National Natural Science Foundation of China (31125027, 31372556), and fund of Xiamen south ocean research center (13GZP002NF08). We thank XinXin You, Yue Feng, GuanXing Chen, RiBei Fu, JinTu Wang, Ying Qiu and Jie Bai (BGI-Shenzhen, China) very much for their hard work in preparing the manuscript and analyses. We thank Jiafu Liu (Fujian Ningde Municipal Station of Fishery Technical Extension) for sample collection. We also thank Qiong Liu and JiaZan Ni (College of Life Sciences, Shenzhen University, China) for help with selenoprotein prediction.

